# MICU1 regulates mitochondrial cristae structure and function independent of the mitochondrial calcium uniporter channel

**DOI:** 10.1101/803213

**Authors:** Dhanendra Tomar, Manfred Thomas, Joanne F. Garbincius, Devin W. Kolmetzky, Oniel Salik, Pooja Jadiya, April C. Carpenter, John W. Elrod

**Author notes:** Correspondence: John W. Elrod, PhD, Center for Translational Medicine, 3500 N Broad St, MERB 949, Philadelphia, PA 19140, LAB, elrodlab.org.

## Abstract

MICU1 is an EF-hand-containing mitochondrial protein that is essential for gating of the mitochondrial Ca^2+^ uniporter channel (mtCU) and is reported to interact directly with the pore-forming subunit, MCU and scaffold EMRE. However, using size-exclusion proteomics, we found that MICU1 exists in mitochondrial complexes lacking MCU. This suggests that MICU1 may have additional cellular functions independent of regulating mitochondrial Ca^2+^ uptake. To discern mtCU-independent MICU1 functions, we employed a proteomic discovery approach using BioID2-mediated proximity-based (<10nm) biotinylation and subsequent LC-MS detection. The expression of a MICU1-BioID2 fusion protein in *MICU1^-/-^* and *MCU^-/-^* cells allowed the identification of total vs. mtCU-independent MICU1 interactors. Bioinformatics identified the Mitochondrial Contact Site and Cristae Organizing System (MICOS) components MIC60 (encoded by the *IMMT* gene) and Coiled-coil-helix-coiled-coil helix domain containing 2 (CHCHD2) as novel MICU1 interactors, independent of the mtCU. We demonstrate that MICU1 is essential for proper proteomic organization of the MICOS complex and that MICU1 ablation results in altered cristae organization and mitochondrial ultrastructure. We hypothesize that MICU1 serves as a MICOS calcium sensor, since perturbing MICU1 is sufficient to modulate cytochrome c release independent of mitochondrial Ca^2+^ uptake across the inner mitochondrial membrane (IMM). Here, we provide the first experimental evidence suggesting that MICU1 regulates cellular functions independent of mitochondrial calcium uptake and may serve as a critical mediator of Ca^2+^-dependent signaling to modulate mitochondrial membrane dynamics and cristae organization.

## Introduction

Calcium (Ca^2+^) is well characterized as an essential second messenger that regulates numerous cellular functions by binding distinct Ca^2+^ sensing domains or motifs present on numerous proteins (Bagur and Hajnoczky, 2017; Carafoli, 2002, 2003). Most Ca^2+^ sensors contain more than one Ca^2+^ binding domain, often with varied affinities for Ca^2+^ binding, resulting in diverse and graded functions in a variety of cellular processes (Bagur and Hajnoczky, 2017; Carafoli, 2002, 2003; Tadross et al., 2008). The Ca^2+^ concentration varies greatly between different cellular compartments, and thus Ca^2+^ sensors are strategically localized for subcellular/organelle specific signaling (Bagur and Hajnoczky, 2017; Rizzuto et al., 2012; Rizzuto and Pozzan, 2006). Mitochondria actively regulate their Ca^2+^ concentration and contain Ca^2+^ sensors to mediate anterograde and retrograde signaling (Bagur and Hajnoczky, 2017; Rizzuto et al., 2012). Examples include Mitochondrial Rho GTPases (MIROs) localized to the outer mitochondrial membrane (OMM), and Mitochondrial calcium uptake proteins (MICUs) localized to the inter-membrane space (IMS) side of the IMM (Bagur and Hajnoczky, 2017; Fransson et al., 2003; Perocchi et al., 2010; Plovanich et al., 2013). MIRO Ca^2+^ sensing is essential for mitochondrial trafficking and structural homeostasis (Bagur and Hajnoczky, 2017; Fransson et al., 2003; Frederick et al., 2004; Nemani et al., 2018; Saotome et al., 2008), while MICUs are known to gate the mitochondrial calcium uniporter channel (mtCU) and regulate its open probability (Csordas et al., 2013; Liu et al., 2016; Mallilankaraman et al., 2012b; Patron et al., 2014; Plovanich et al., 2013).

The mtCU is a highly selective Ca^2+^ channel necessary for acute Ca^2+^ entry to the mitochondrial matrix (Baughman et al., 2011; De Stefani et al., 2011; Kirichok et al., 2004; Rizzuto et al., 2012). The mtCU consists of multiple subunits, namely the pore-forming component Mitochondrial Calcium Uniporter (MCU) and its homolog, MCUB; the regulatory scaffolds MCU Regulator 1 (MCUR1) and Essential MCU Regulator Element (EMRE); and the Ca^2+^ sensors Mitochondrial Calcium Uptake proteins 1, 2, and 3 (MICU1, MICU2 and MICU3) (Baughman et al., 2011; De Stefani et al., 2011; Mallilankaraman et al., 2012a; Perocchi et al., 2010; Plovanich et al., 2013; Raffaello et al., 2013; Sancak et al., 2013; Tomar et al., 2016). MICU1 is essential to mtCU regulation, by directly binding MCU and EMRE and its expression correlates with tissue-dependent differences in mitochondrial calcium uptake (Csordas et al., 2013; Mallilankaraman et al., 2012b; Paillard et al., 2018; Patron et al., 2014; Perocchi et al., 2010; Phillips et al., 2019; Plovanich et al., 2013; Sancak et al., 2013; Xing et al., 2019).

Loss-of-function mutations in *MICU1* induce proximal myopathy, learning difficulties, movement disorder, fatigue, and lethargy in humans (Lewis-Smith et al., 2016; Logan et al., 2014) and deletion of *Micu1* in mouse models causes perinatal lethality (Antony et al., 2016; Liu et al., 2016). Recently, genetic mutants were generated to characterize mtCU regulation in Drosophila (Tufi et al., 2019). Intriguingly, Tufi et al. reported that a *MICU1* loss-of-function mutation resulted in Drosophila lethality, which could not be rescued by a concurrent *MCU* loss-of-function mutation that completely ablated mitochondrial Ca^2+^ (_m_Ca^2+^) uptake and subsequent mitochondrial permeability transition pore opening (Tufi et al., 2019). This observation suggests that the lethal phenotype of MICU1-null flies was not solely a result of aberrant mtCU-dependent Ca^2+^ uptake. This raises the possibility that MICU1 has mtCU-independent functions, which are vital for mitochondrial function and survival. Indeed, MICU1 knockout models show distinct abnormalities in mitochondrial ultrastructure that are not observed in any other mtCU knockout models (Bick et al., 2017; Liu et al., 2016; Luongo et al., 2015; Tomar et al., 2016). Additionally, MICU1 protein is reported to have high mobility within the IMM as compared to the MCU (Hoffman et al., 2013), suggesting that MICU1 could be associated with other complexes in the mitochondria. These observations led us to hypothesize that MICU1 regulates other essential mitochondrial processes beyond calcium uptake.

To discover mtCU-independent functions of the MICU1, we utilized a proximity-based biotinylation approach by generating a MICU1-BioID2 fusion protein. BioID2 is a recently developed, highly-efficient promiscuous biotin ligase which enables the detection of protein-protein interactions in living cells (Kim et al., 2016). We reconstituted *MICU1*^-/-^ cells with MICU1-BioID2-HA to avoid aberrant localization associated with overexpression to characterize the entire MICU1 interactome. We also expressed MICU1-BioID2-HA in *MCU*^-/-^ cells to define mtCU-independent interactions. Through a comparative analysis of mass-spectrometry, we identified proteins whose interaction with MICU1 was unaffected by the loss of MCU. Here, we report that MICU1 directly interacts with the Mitochondrial Contact Site and Cristae Organizing System (MICOS) components MIC60, and CHCHD2 in an MCU-independent manner. Our results suggest that MICU1 confers calcium sensing to the MICOS for cell signaling-dependent changes in cristae structure and function.

## Results and Discussion

### Observation of MICU1 localization independent of the mtCU

To define the mtCU-independent molecular functions of the MICU1, we first utilized size exclusion chromatography to characterize the native organization of MICU1-containing protein complexes. Total cell lysates prepared from HEK293T *MCU*^+/+^ and *MCU*^-/-^ cells were fractioned in non-reducing conditions by Fast Protein Liquid Chromatography (FPLC). FPLC fractions ranging from ∼10kDa to ∼900kDa were collected, concentrated and examined for the MICU1 protein complexes using reducing SDS-PAGE and Western blotting (Figure 1A, 1B). MICU1 forms distinct high-molecular weight (MW) protein complexes ranging from ∼200 kDa to ∼700 kDa (Figure 1A, 1B). Intriguingly, the loss of MCU does not have a substantial effect on the overall distribution of MICU1-containing high-molecular weight (MW) protein complexes (Figure 1A, 1B). Next, we examined mitochondrial sub-localization of MCU and MICU1 by immunofluorescent detection of native MCU and FLAG-tagged MICU1 in *Micu1*^-/-^ mouse embryonic fibroblasts (MEFs) to enable accurate detection of MICU1 and avoid aberrant localization associated with overexpression (Figure 1C). The deletion of MICU1 in MEFs was confirmed by Western blotting (Figure S1A). Line-scan analysis of the mitochondrial network clearly shows that MICU1 co-localizes with MCU, but also distributes to sub-mitochondrial regions lacking MCU (Figure 1C, 1D). These results suggest that MICU1 is present in mitochondrial protein complexes where the mtCU is absent.

**Figure 1.**
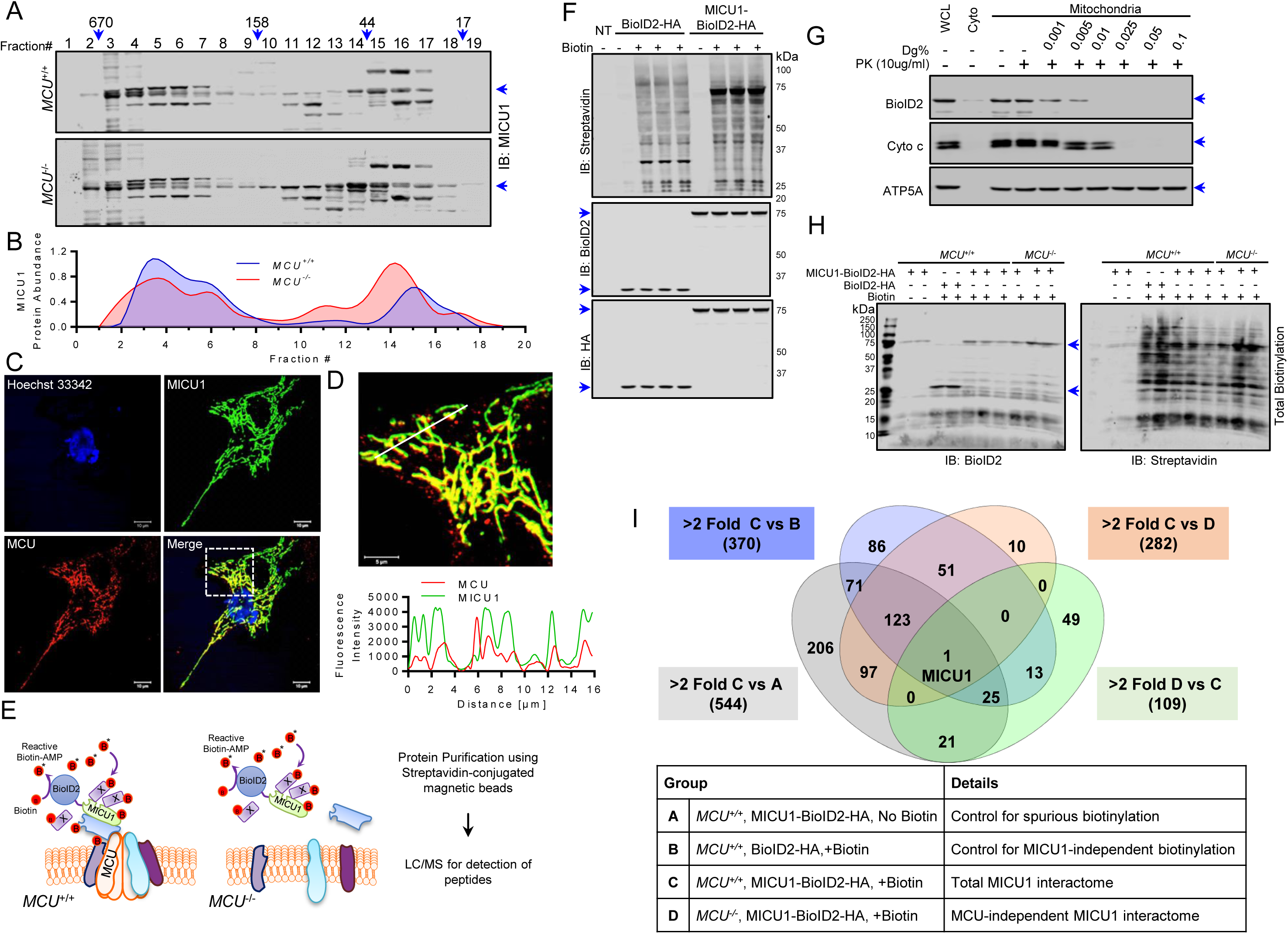
Identification of mtCU independent MICU1 interactors. **(A)** Cleared cell lysates isolated from WT and *MCU*^-/-^ HEK293T cells were fractionated by FPLC-based size exclusion chromatography. The protein fractions ranging from ∼10kDa to ∼900kDa were collected, concentrated, and subjected to immunoblotting using the MICU1 antibody. **(B)** Densitometry was performed to generate chromatographs of the MICU1 distribution in different fractions in Figure 1A. **(C-D)** MEFs stably expressing MICU1-FLAG were cultured on collagen-coated cover glass, were fixed with 4%PFA, and co-immunofluorescence was performed using FLAG and MCU antibodies. Images were acquired using an LSM 510 META Laser Scanning Microscope (Carl Zeiss, Inc.). Scale bar C= 10µm, C inset= 5µm, D= 5µm. **(E)** Experimental outline for the utilization of MICU1-BioID2-HA fusion protein to identify the mtCU-independent MICU1 interactors via biotin-based proximity labeling. **(F)** HEK293T *MICU1*^-/-^ cells were transfected with plasmids encoding BioID2-HA, and MICU1-BioID2-HA. Cells were cultured in the presence of biotin (50µM) for 16h to induce the biotinylation. Cells were collected, washed twice with PBS, lysed in BioID2 lysis buffer, western blotting with the indicated antibodies. **(G)** Mitochondria were isolated from HEK293T *MICU1*^-/-^ cells reconstituted with MICU1-BioID2-HA. Mitochondrial fractions were subjected to increasing digitonin concentrations to permeabilize the outer mitochondrial membrane (OMM) and inner mitochondrial membrane (IMM). Proteinase K treatment was performed to cleave the exposed proteins, and mitochondrial fractions were probed with indicated antibodies. **(H)** HEK293T *MICU1*^-/-^ and HEK293T *MCU*^-/-^ cells were transfected with plasmids encoding BioID2-HA or MICU1-BioID2-HA. Cells were cultured in the presence or absence of biotin (50µM) for 16h. Cells were collected, washed with PBS 2 times, and lysed in BioID2 lysis buffer. An aliquot of the lysates were subjected to western blotting using the anti-BioID2 antibody and Streptavidin to validate effective biotinylation of cellular proteins. **(I)** Protein samples were subjected to Streptavidin based pull-down and digested with trypsin. LC-MS/MS analysis of 10% of total digests in duplicate runs was performed by on-line analysis of peptides by a high-resolution, high-accuracy LC-MS/MS system (Thermo Fisher Scientific). Estimated protein abundance after global sample normalization was used to compare different groups.

Next, we characterized the mtCU-independent interactome of MICU1. We generated a MICU1-BioID2-HA fusion protein to enable the biotinylation of interactors (<10nm) in *MCU*^+/+^ and *MCU*^-/-^ cells to distinguish the mtCU-dependent vs. -independent MICU1 interactomes (Figure 1E). Expression, biotin ligase activity, sub-mitochondrial localization and reconstitution of _m_Ca^2+^ uptake regulation of MICU1-BioID2-HA fusion protein was confirmed in *MICU1^-/-^* cells expressing the MICU1-BioID2-HA fusion protein (Figure 1F, 1G, S1B). These data show that our fusion construct was properly localized and that mtCU-dependent calcium uptake was not altered in our discovery system. Next, we expressed the MICU1-BioID2 or BioID2 control in HEK293T *MICU1^-/-^* cells (hereafter *MCU^+/+^*) and HEK293T *MCU^-/-^* cells (hereafter *MCU^-/-^*) (Figure 1H). Biotinylation of proteins proximal to MICU1-BioID2 was induced by culturing cells in presence of biotin (50µM) for 16h (Figure 1H). MICU1-BioID2-HA protein expression and biotin ligase activity were confirmed via Western blotting (Figure 1H). Next, biotinylated proteins were purified from cell lysates using streptavidin-conjugated magnetic beads. Peptides were generated from the purified biotinylated proteins by tryptic digestion, and LC-MS was performed (Kim et al., 2016). Comparative analysis of MICU1 proximal proteins identified in *MCU*^+/+^ vs. *MCU*^-/-^ cells was performed (Figure 1I). The MICOS components MIC60, CHCHD3, CHCHD2, APOO, and APOOL emerged as ‘hits’ from a single multiprotein complex present at the inner mitochondrial membrane (IMM), and proximity to MICU1 was unaltered in *MCU*^-/-^ cells (Figure 1I, Table S1).

### MICU1 directly interacts with MIC60 and CHCHD2 in the MICOS complex

The three core MICOS components along with OPA1, which is also involved in cristae organization, emerged as MICU1 proximal proteins in our proteomic screen (Figure 1I, Table S1). However, the loss of MCU results in loss of the MICU1:OPA1 interaction, while the MICU1 interaction with the core MICOS components is preserved (Figure 1I, Table S1). This observation suggests that MICU1 could be an integral component of the MICOS complex and involved in mitochondrial cristae organization, independent of the mtCU and mitochondrial calcium uptake. To assess if MICU1 directly binds MIC60, CHCHD2, and CHCHD3 we co-expressed MIC60-FLAG, CHCHD2-FLAG, or CHCHD3-FLAG with MICU1-HA and 48h post-transfection, we performed immunoprecipitation (IP) with FLAG-conjugated magnetic beads. IP’d products were analyzed by SDS-PAGE and Western blotting for HA immunoreactivity, to detect MICU1, and FLAG expression, to detect MICOS components (Figure 2A). To control for level of expression we blotted for FLAG-tagged MIC60 (∼90kDa), CHCHD2 (∼16kDa), and CHCHD3 (∼26kDa) and HA-tagged MICU1 (∼55kDa) (Figure 2A). All were expressed to similar degrees, but only MIC60 and CHCHD2 pulled-down with MICU1 (Figure 2A). We validated this result by reverse IPs and confirmed the interaction of MICU1-FLAG with endogenous MIC60 and CHCHD2, but not with CHCHD3 (Figure S2A). This suggests that MICU1 may directly interact with MIC60 and CHCHD2, but not with CHCHD3 and that its biotinylation in our discovery assay was likely due to its general proximity to MICU1. To substantiate the MICU1 interaction with MIC60 and CHCHD2 we performed co-immunofluorescence labeling and imaged to examine sub-mitochondrial localization (Figure 2B, 2C, 2D, 2E). The line-scan profile shows distinct pixels with spectral overlap of MICU1 with MIC60, and MICU1 with CHCHD2 (Figure 2C, 2E). Together, these data suggest that MICU1 directly interacts with the core MICOS components.

**Figure 2.**
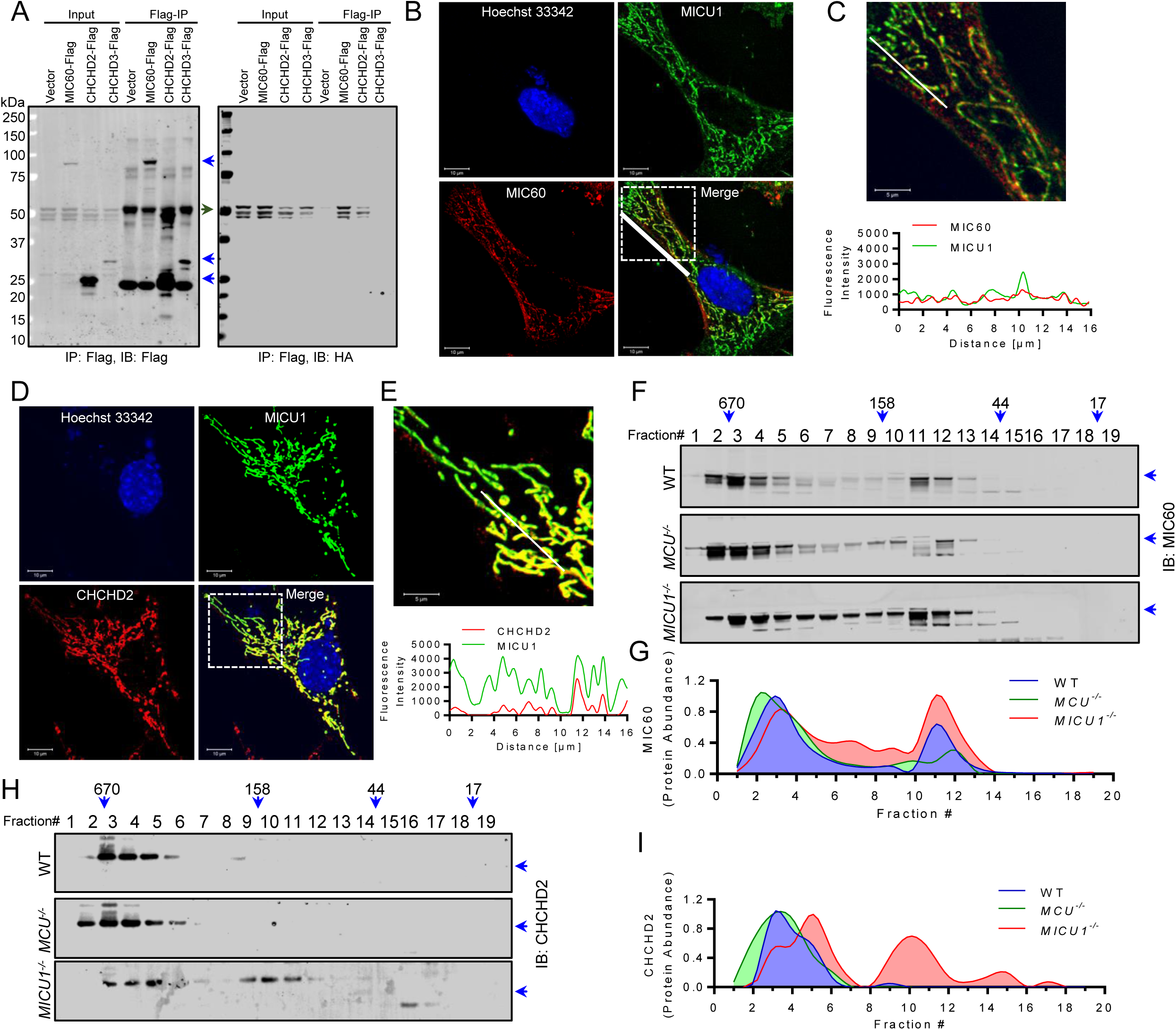
MICU1 directly interacts with MICOS components and is essential for the formation of high-molecular-weight MICOS. **(A)** MICU1-HA and FLAG-tagged MICOS components co-expressed in the HEK293T *MICU1*^-/-^ cells. Cell lysates of equal protein content were subjected to FLAG-immunoprecipitation (IP), and IP products were probed with FLAG and HA antibodies to detect the interaction between MICU1 and MICOS components. **(B-E)** MICU1-FLAG expressing MEFs were cultured on the collagen-coated cover glass and fixed with PFA, then co-immunofluorescence was performed using FLAG and MIC60 (B, C), or CHCHD2 (D, E) antibodies. Images were acquired using an LSM 510 META Laser Scanning Microscope (Carl Zeiss, Inc.). Merged images show the co-localization between the MICU1 and MIC60 or CHCHD2. Line scan shows the presence of MICU1 with both the MICOS components. Scale bar B, D= 10µm; C, E inset= 5µm. **(F, H)** FPLC was performed using cleared cell lysates from WT, *MCU*^-/-^, and *MICU1*^-/-^ HEK293T cells, and fractions were collected and subjected to western blotting using the MIC60 (F), and CHCHD2 (H) antibodies. The loss of MICU1 alters the size distribution of the multimeric MICOS complex. **(G, I)** Densitometry was performed to generate chromatographs of the MICOS components distribution in different molecular weight fractions in Figure 2F and 2H.

To further characterize the functional relevance of MICU1 interaction with MICOS components, we performed FPLC to fractionate the high-MW MICOS complex in WT, *MCU*^-/-,^ and *MICU1*^-/-^ cells. Immunoblots of 19 fractions ranging from ∼10kDa to ∼900kDa were probed for MIC60, CHCHD2 and CHCHD3 expression; all showed robust immunoreactivity in native protein complexes ranging from ∼400-700 kD (Figure 2F-2I, S2B, S2C). Interestingly, genetic deletion of *MCU* had no effect on the overall size or fraction distribution of the multi-subunit MICOS complex (Figure 2F-2I, S2B, S2C). However, the loss of MICU1 expression resulted in a rightward shift, decrease in overall MW, of MIC60, CHCHD2, and CHCHD3 containing complexes (Figure 2F-2I, S2B, S2C) suggesting that MICU1 may play an integral role in MICOS complex assembly or stability.

### MICU1 is essential for the maintenance of mitochondrial ultrastructure and cristae organization

The MICOS is essential for maintenance of mitochondrial membrane topology and bottleneck formation (Friedman et al., 2015; Harner et al., 2011; Tarasenko et al., 2017; van der Laan et al., 2016). The MICOS is localized at the intersection of the IMM and OMM, which results in the formation of membrane contact sites at cristae junctions (Friedman et al., 2015; Harner et al., 2011; Tarasenko et al., 2017; van der Laan et al., 2016). Ca^2+^ is reported to modulate the cristae structure (Gottschalk et al., 2018; Greenawalt et al., 1964), however, no Ca^2+^-sensing protein has yet been identified as an essential component of the MICOS. To discern if MICU1 serves as a conduit for calcium-dependent regulation of the MICOS, we examined if genetic loss of *MICU1* had any effect on mitochondrial ultrastructure and cristae junctions. In agreement with previous reports, transmission electron microscopy (TEM) revealed gross changes in mitochondrial ultrastructure of cells lacking MICU1 (Figure 3A-3D). Careful quantitative analysis of TEM images showed that mitochondrial perimeter, mitochondrial Feret diameter (the distance between the two parallel planes restricting the object perpendicular to that direction), and aspect ratio are significantly reduced in *MICU1*^-/-^ cells (Figure 3B-3D). This suggests that mitochondria are less filamentous in *MICU1*^-/-^ cells. Next, we analyzed the inter-cristae junction (distance between cristae) and the cristae junction width (distance between IMM of the same cristae) in WT and *MICU1*^-/-^ cells. The inter-cristae junction distance is reported to be directly proportional to cristae density. *MICU1*^-/-^ cells displayed a significant increase in both the inter-cristae junction distance and cristae junction width, as compared to WT cells (Figure 3E, 3F).

**Figure 3.**
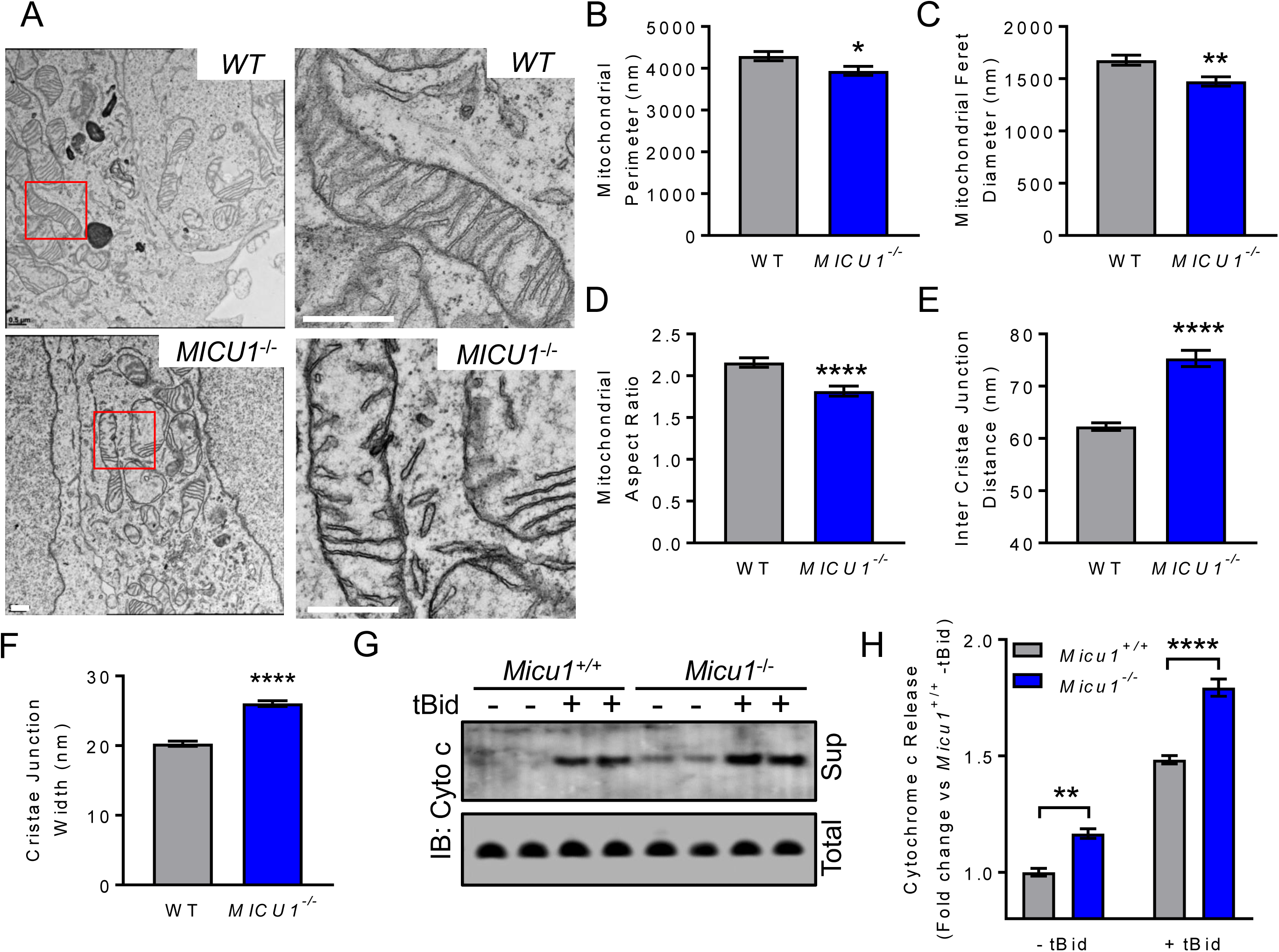
*MICU1*^-/-^ cells show altered cristae structure and enhanced cytochrome c release. **(A)** WT and *MICU1*^-/-^ HEK293T cells were grown on Thermanox® Cover Slips and processed for the TEM imaging. Images were acquired by a Zeiss LIBRA120 TEM equipped with Gatan UltraScan, 1000 2k x 2k CCD EFTEM, energy filtering. *MICU1*^-/-^ cells show distinct alterations in cristae organization and mitochondrial ultrastructure. Scale bar = 500nm. **(B-F)** TEM images were analyzed and quantitated using the Image J Fiji. Mitochondrial perimeter (B), feret diameter (C), aspect ratio (D), inter-cristae junction distance (E), and cristae junction width (F) were plotted. Statistical significance was determined using t-test. * indicates p<0.05. n=200-300 mitochondria. **(G)** *Micu1*^+/+^ and *Micu1*^-/-^ MEFs were grown in 150mm^2^ culture dishes, and cytochrome c release assay was performed. *Micu1*^-/-^ MEFs show enhanced basal and tBid-induced cytochrome c release. **(H)** Densitometry was performed to quantify the cytochrome c release in different groups in Figure 3G. Statistical significance was determined using t-test. * indicates p<0.05. n=3.

The bottleneck structure of cristae is essential to the maintenance of the mitochondrial respiratory chain complexes (Friedman et al., 2015; van der Laan et al., 2016). Disorganization and cristae remodeling is associated with the release of cytochrome c from bottlenecks, which subsequently induces cell death signaling pathways (Scorrano et al., 2002). To further define the role of MICU1 in cristae regulation, we monitored tBid-induced cytochrome c release in primary *Micu1*^-/-^ mouse embryonic fibroblasts (MEFs). The loss of MICU1 resulted in increased basal cytochrome c release and this was potentiated after tBID treatment (Figure 3G, 3H). To rule out possible indirect effects of MICU1 regulation of mtCU-Ca^2+^ uptake on cristae structure, we examined _m_Ca^2+^ uptake in *Chchd2*^-/-^ MEFs. CHCHD2 was previously identified as a core MICOS component and its genetic deletion results in abnormal cristae organization (Meng et al., 2017). *WT* and *Chchd2*^-/-^ MEFs were permeabilized with digitonin in the presence of thapsigargin to monitor _m_Ca^2+^ uptake independent of plasma membrane and ER Ca^2+^ transport using ratiometric Ca^2+^ sensor. *Chchd2*^-/-^ cells showed no change in _m_Ca^2+^ uptake suggesting that altered cristae structure alone is insufficient to impact mtCU-dependent _m_Ca^2+^ uptake (Figure S3A, S3B). Further, we found that loss of CHCHD2 had no effect on mitochondrial calcium efflux (rate of Ca^2+^ exiting the matrix after Ru360 blockade of MCU; Figure S3A, S3C). We also examined the high-MW/functional mtCU complex in *Chchd2*^-/-^ MEFs by FPLC-based protein fractionation and observed no change in MCU distribution (Figure S3D, S3E). These observations suggest that altered cristae structure alone does not have a significant impact on mtCU assembly or _m_Ca^2+^ dynamics.

In summary, we characterized the MICU1 interactome and identified a distinct involvement in cristae organization independent of the mtCU and mitochondrial calcium uptake. Our study reveals a direct interaction between MICU1 and core MICOS components and shows that this interaction is essential to form the functional MICOS complex. This mechanism could explain the lethal phenotype observed in MICU1 knockout models (Antony et al., 2016; Liu et al., 2016; Tufi et al., 2019), as our results suggest that loss of MICU1 could induce both necrotic, as well as apoptotic, death signaling events independent of matrix Ca^2+^ overload. Our results also highlight the need to reappraise the MICU1/mtCU literature as some of the reported phenotypes may be influenced by alterations in the function/structure of the MICOS, rather than dependent on changes in mitochondrial calcium uptake. Further research is needed to define the precise interaction of MICU1 with MICOS components to hopefully identify tools to enable the dissection of mtCU-dependent vs. independent functions in mitochondrial biology. In summary, the current study identified a novel role for MICU1 in regulating the cristae junction, independent of the mtCU, which is essential for mitochondrial physiology.

## Materials and Methods

Key resources table.

**Key resources table**
| Reagent type or resource | Designation | Source or reference | Identifiers | Additional information |
| --- | --- | --- | --- | --- |
| Cell lines | HEK293T WT | (Sancak et al., 2013) |  |  |
|  | HEK293T *MCU*<sup>-/-</sup> | (Sancak et al., 2013) |  |  |
|  | HEK293T *MICU1*<sup>-/-</sup> | (Sancak et al., 2013) |  |  |
|  | MEF *Micu1*<sup>+/+</sup> | This study |  |  |
|  | MEF *Micu1*<sup>-/-</sup> | This study |  |  |
|  | MEF *Chchd2*<sup>+/+</sup> | (Meng et al., 2017) |  |  |
|  | MEF *Chchd2*<sup>-/-</sup> | (Meng et al., 2017) |  |  |
| Plasmids | MCS-BioID2-HA | Addgene (Kim et al., 2016) | Cat# 74224 |  |
|  | Start-BioID2-HA | This study |  |  |
|  | MICU1-BioID2-HA | This study |  |  |
|  | hMICU1-Myc-DDK | OriGene Technologies | Cat# RC200921 |  |
|  | MICU1-FLAG | OriGene Technologies | Cat# MR207652 |  |
|  | pCMV6-AC-HA | OriGene Technologies | Cat# PS100004 |  |
|  | MICU1-HA | This study |  |  |
|  | MIC60-FLAG | OriGene Technologies | Cat# MR216091 |  |
|  | CHCHD3-FLAG | OriGene Technologies | Cat# MR202692 |  |
|  | CHCHD2-FLAG | OriGene Technologies | Cat# MR223513 |  |
| Antibodies | Anti-MIC60<br>IMMT Rabbit<br>Polyclonal Antibody<br>(10179-1-AP) | Proteintech | Cat# 10179-1-AP | WB 1:1000<br>IF 1:200<br>PLA 1:200 |
|  | Anti-CHCHD3<br>Rabbit Polyclonal<br>Antibody | Proteintech | Cat# 25625-1-AP | WB 1:1000 |
|  | Anti-CHCHD2 Rabbit<br>Polyclonal Antibody | Proteintech | Cat# 19424-1-AP | WB 1:1000<br>IF 1:200<br>PLA 1:200 |
|  | Anti-MCU<br>Rabbit Polyclonal<br>Antibody | Sigma-Aldrich | Cat#<br>HPA016480-100UL | WB 1:1000<br>IF 1:200<br>PLA 1:200 |
|  | Monoclonal ANTI-<br>FLAG® M2 antibody | Sigma-Aldrich | Cat# F1804-1MG | WB 1:2000<br>IF 1:200<br>PLA 1:200 |
|  | Anti-HA High Affinity,<br>from rat IgG1 | Sigma-Aldrich | Cat#<br>11867423001 | WB 1:3000 |
|  | Anti-MICU1<br>Rabbit Polyclonal<br>Antibody | In-house (Tomar et al.,<br>2016) |  | WB 1:500 |
|  | Anti-BioID2 Chicken<br>Polyclonal Antibody | BioFront Technologies | Cat# BID2-CP-100 | WB 1:500 |
|  | Cytochrome c<br>Antibody (A-8) Alexa<br>Fluor® 680 | Santa Cruz Biotechnology | Cat# sc-13156<br>AF680 | WB 1:5000 |
|  | IRDye 800CW<br>Streptavidin | LI-COR Biosciences | Cat# 925-32230 | WB 1:10000 |
|  | IRDye® 680RD Goat<br>anti-Mouse IgG (H +<br>L) | LI-COR Biosciences | Cat# 925-68070 | WB 1:10000 |
|  | IRDye® 680RD Goat anti-Rat IgG (H + L) | LI-COR Biosciences | Cat# 925-68076 | WB 1:10000 |
|  | IRDye® 680RD Goat anti-Rabbit IgG (H + L) | LI-COR Biosciences | Cat# 926-68071 | WB 1:10000 |
|  | Goat Anti-Chicken IgY H&L (Alexa Fluor® 680) | Abcam | Cat# ab175779 | WB 1:10000 |
|  | IR Dye Goat anti Rabbit 800CW | LI-COR Biosciences | Cat# 926-32211 | WB 1:10000 |
|  | IRDye® 800CW Goat anti-Mouse IgG (H + L) | LI-COR Biosciences | Cat# 926-32210 | WB 1:10000 |
|  | Donkey anti-Mouse IgG (H+L) Highly Cross-Adsorbed Secondary Antibody, Alexa Fluor 488 | Thermo Fisher Scientific | Cat# A-21202 | IF 1:500 |
|  | Goat anti-Rabbit IgG (H+L) Highly Cross-Adsorbed Secondary Antibody, Alexa Fluor Plus 647 | Thermo Fisher Scientific | Cat# A32733 | IF 1:500 |
| Magnetic Beads | ANTI-FLAG(R) M2 MAGNETIC BEADS | Sigma-Aldrich | Cat# M8823-1ML |  |
|  | DYNAL MyOne Dynabeads | Thermo Fisher Scientific | Cat# 65-001 |  |
|  | Streptavidin C1 |  |  |  |
| Commercial assay or kit | Duolink® In Situ Red Starter Kit Mouse/Rabbit | Sigma-Aldrich | Cat# DUO92101-1KT | PLA |
|  | Recombinant Human BID (Caspase-8-cleaved) Protein, CF | R&D Systems | Cat# 882-B8-050 | Cytochrome c release assay |
| Staining reagents | Fura-FF | AAT Bioquest | Cat# 21028 | Ca<sup>2+</sup> detection |
|  | Hoechst 33342 | Thermo Fisher Scientific | Cat# 62249 | Nucleus Staining |
| Software, algorithm | GraphPad Prism | https://www.graphpad.com/ |  |  |
|  | Image J Fiji | https://imagej.net/Fiji |  |  |
|  | Zeiss Zen | https://www.zeiss.com |  |  |

### Plasmids Construction

To generate Start-BioID2-HA plasmid, BioID2 was PCR-amplified from the MCS-BioID2-HA plasmid using primers designed to introduce an ATG start codon immediately downstream of the BamHI restriction site of the MCS. The PCR product was cloned via BamHI and HindIII into the MCS-BioID2-HA plasmid (Addgene #74224). To generate the MICU1-BioID2-HA, MICU1 was PCR amplified from the hMICU1-Myc-DDK plasmid using primers to introduce a 5’ AgeI and a 3’ BamHI restriction site. The PCR product was cloned via AgeI and BamHI into the MCS-BioID2-HA plasmid (Addgene #74224). MICU1-HA plasmid was generated by cleaving the MICU1 fragment from the MICU1-FLAG plasmid using the SgfI-MluI restriction sites and inserted at the same sites in pCMV6-AC-HA vector. All plasmids were confirmed using the restriction digestion and DNA sequencing.

### Cell culture

HEK293T WT, HEK293T *MCU*^-/-^ and HEK293T *MICU1*^-/-^ cells were grown in Dulbecco’s Modification of Eagle’s Medium with 4.5 g/L glucose, L-glut, and Na Pyr medium (Corning Cellgro, Cat#10-013-CV) supplemented with 10% fetal bovine serum (Peak Serum, Cat#PS-FB3), 1% penicillin/streptomycin (Sigma-Aldrich, Cat# P0781-100ML) at 37°C in the presence of 5% CO_2_. Mouse embryonic fibroblasts isolated from the *Micu1*^fl/fl^ mouse were immortalized by infecting the cells with SV40 large T antigen-expressing adenovirus. The immortalized *Micu1*^fl/fl^ MEFs serve as *Micu1*^+/+^ control cells. *Micu1*^-/-^ MEFs were generated by transducing the *Micu1*^+/+^ MEFs with adenovirus encoding Cre-recombinase (Ad-Cre). MEFs were grown in Dulbecco’s Modification of Eagle’s Medium with 4.5 g/L glucose, L-glut, and Na Pyr medium (Corning Cellgro, Cat#10-013-CV) supplemented with 10% fetal bovine serum (Peak Serum, Cat#PS-FB3), 1% Gibco® MEM Non-Essential Amino Acids (Thermo Fisher Scientific, Cat# 11-140-050), 1% penicillin/streptomycin (Sigma-Aldrich, Cat# P0781-100ML), at 37°C in the presence of 5% CO_2_. The *Chchd2*^+/+^ and *Chchd2*^-/-^ MEFs were cultured as described earlier (Meng et al., 2017). To exogenously express MICU1, MIC60, CHCHD3, and CHCHD2, HEK293T cells were transfected with the Fugene HD transfection reagent (Promega, Cat#E2311) as per manufacturer instruction. To generate the MEFs stably expressing MICU1-FLAG, immortalized WT MEFs were transfected with MICU1-FLAG plasmid (OriGene Technologies, Cat#MR207652) using the Fugene HD transfection reagent (Promega, Cat#E2311). 24h post-transfection, culture media was replaced with media supplemented with the 500 µg/mL G418 (Thermo Fisher Scientific, Cat#10131035). Fresh culture media supplemented with G418 was replaced at two-day interval, until all the dying cells were cleared. After incubation for two weeks, the cells were maintained in DMEM supplemented with 200 µg/mL G418.The protein expression was validated by western blotting and immunofluorescence.

### Immunoblotting

Cells were harvested, washed with ice-cold PBS and lysed in 1X RIPA lysis buffer (EMD Millipore, Cat#20-188) supplemented with SIGMAFAST™ Protease Inhibitor Cocktail (Sigma-Aldrich, Cat#S8830). Protein concentrations were determined by Pierce 660nm Protein Assay (Thermo Fisher Scientific, Cat#22660) and equal ug of protein were separated by electrophoresis on NuPAGE 4-12% Bis-Tris protein gel (Thermo Fisher Scientific, Cat#WG1402BOX), in denaturing conditions. Protein was electroblotted on PVDF membrane (EMD Millipore, Cat#IPFL00010). Following the transfer, the membrane was incubated in Blocking Buffer (Rockland, Cat#MB-070) for 1h at room temperature, followed by overnight incubation with specific primary antibody at 4°C. After incubation the membrane was washed with TBS-T (TBS containing 0.1% Tween 20) 3 times for 10 min each and then incubated with specific secondary antibody for 1h at room temperature. The membrane was similarly washed 3 times with TBS-T and then imaged on an LI-COR Odyssey system.

### Sub-mitochondrial protein localization assay

Mitochondria were isolated as described earlier (Tomar et al., 2015). Briefly, cells were grown in 150 mm^2^ culture dishes, washed with PBS, and resuspended in isotonic mitochondria isolation buffer (10 mM HEPES, pH 7.5, containing 200 mM mannitol, 70 mM sucrose, and 1 mM EGTA). The cell suspension was homogenized by Dounce homogenizer, centrifuged at 500 g for 10 min at 4°C. Supernatant was collected and centrifuged at 12,000 g for 15 min at 4°C to obtain crude mitochondrial pellet. The pellet was resuspended in mitochondria isolation buffer and washed 2 times using the centrifuge at 12,000 g for 15 min at 4°C. The mitochondrial pellet was resuspended in intracellular buffer (120 mM KCl, 10 mM NaCl, 1 mM KH2PO4, 20 mM HEPES-Tris, pH 7.2) and permeabilized with varying digitonin concentrations and digested with proteinase K (10 µg/mL) for 10min at room temperature. Proteinase K digestion was stopped by adding SIGMAFAST™ Protease Inhibitor Cocktail (Sigma-Aldrich, Cat#S8830) and 2X SDS-loading dye and heating the samples at 95°C for 10min. The immunoblotting was performed using specified antibodies.

### Biotinylation and mass spectrometry analysis

To induce BioID2-mediated protein biotinylation, cells were cultured with media supplemented with 50 μM biotin for 16h. Cells were collected, washed with PBS 2 times, and lysed in BioID2 lysis buffer (50 mM Tris, pH 7.4, 500 mM NaCl, 2% Triton X-100, 0.4% SDS, 1 mM dithiothreitol) supplemented with SIGMAFAST™ Protease Inhibitor Cocktail (Sigma-Aldrich, Cat#S8830). The cell suspension was sonicated for 2 times each for 1 min at an output level of 40 (Vibra-Cell, Sonics). An equal volume of 50 mM Tris, pH 7.4, was added and the suspension was cleared using centrifugation at 16,500 g for 20 min. The supernatant was used for immunoblotting or Streptavidin based pull-down using MyOne Dynabeads Streptavidin C1. Mass spectroscopy for identification of the biotinylated proteinss was performed as described earlier (Kim et al., 2016).

### FPLC and protein fractionation

Size-exclusion gel filtration was used to separate the high-molecular-weight protein complexes using the fast protein liquid chromatography (ÄKTA Pure FPLC; GE Healthcare) (Tomar et al., 2016). The PBS-equilibrated Superdex 200 10/300 column (GE Healthcare, Cat#17517501) was calibrated with a gel filtration calibration standard (Bio-Rad, Cat#1511901). The cleared cell lysates were directly loaded onto FPLC and fractioned at a flow rate of 0.5 mL/min. Protein fractions were collected in 0.5 mL PBS, concentrated to 50µL volume using an AMICON Ultra-0.5 Centrifugal Filter Devices (with a 3,000 kD cutoff) (EMD Millipore, Cat#UFC500396). Concentrated fractions were then immunoblotted using specific antibodies.

### Co-immunoprecipitation

To study protein-protein interactions, immunoprecipitation experiments were performed as described earlier (Tomar et al., 2016). Briefly, HEK293T cells were co-transfected with the indicated plasmids. After 36h of transfection, cells were harvested, washed with ice-cold PBS and lysed in 1X RIPA lysis buffer (EMD Millipore, Cat#20-188) supplemented with SIGMAFAST™ Protease Inhibitor Cocktail (Sigma-Aldrich, Cat#S8830). Protein concentrations were determined by Pierce 660nm Protein Assay (Thermo Fisher Scientific, Cat#22660) and equal proteins amounts were used for co-immunoprecipitation. Cleared cell lysate was incubated with Anti-FLAG M2 Magnetic Beads (Sigma-Aldrich) on a roller shaker overnight at 4°C. Beads were washed 3 times with RIPA buffer and 2 times with TBS-T, resuspended in 2X SDS-PAGE sample buffers, and then immunoblotting was performed using specific antibodies.

### Co-immunofluorescence

The mitochondrial localization of mtCU, MICOS, and MICU1 was also analyzed by immunofluorescence using standard protocol (Tomar et al., 2015). Briefly, the MEFs stably expressing MICU1-FLAG were grown on collagen-coated 35-mm dishes. Cells were washed with PBS, fixed for 20min with 4% paraformaldehyde, then permeabilized for 15min by 0.15% Triton X-100. Permeabilized cells were blocked using 10% BSA for 45min at room temperature and incubated with primary antibodies overnight at 4°C. After incubation, cells were washed 3 times with blocking reagent and incubated with Alexa Fluor-tagged secondary antibodies for 1h at room temperature. Cells were washed 3 times with PBS, and confocal images were obtained using an LSM 510 META Laser Scanning Microscope (Carl Zeiss, Inc.) at 488- and 647-nm excitations using a 63x oil objective. Images were analyzed and quantitated using ZEN 2010 software (Carl Zeiss, Inc.), and Image J Fiji.

### Proximity ligation assay

Proximity ligation assay (PLA) was used to detect *in situ* MICU1: MICOS interactions. MICU1-FLAG expressing MEFs were seeded on collagen-coated 35-mm dishes. Cells were washed with PBS, fixed for 20min with 4% paraformaldehyde, then permeabilized with 0.15% Triton X-100. PLA was performed as per manufacturer’s instructions (Sigma-Aldrich, Duolink® In Situ Red Mouse/Rabbit Assay). Images were acquired using an LSM 510 META Laser Scanning Microscope (Carl Zeiss, Inc.) using a 63x oil objective.

### Transmission electron microscopy

Transmission electron microscopy (TEM) was utilized to evaluate mitochondrial ultrastructure and cristae organization. HEK293T cells of the indicated genotypes were grown to 80% confluency on 25 mm diameter Thermanox® Cover Slips (Thermo Fisher Scientific, Cat#174985PK) in 6-well plates. Culture media was removed, and cells were fixed with freshly prepared TEM fixation buffer (2% glutaraldehyde, 2% paraformaldehyde in 0.1M sodium cacodylate buffer) for 30 min at room temperature. Fixative was replaced with 0.1M sodium cacodylate buffer, and then samples were processed for TEM imaging. Images were obtained using Zeiss LIBRA120 TEM equipped with Gatan UltraScan, 1000 2k x 2k CCD EFTEM, energy filtering. Images were analyzed and quantitated Image J Fiji.

### Cytochrome c release assay

The cytochrome c release assay was performed as described earlier (Tomar et al., 2015) with slight modifications. Briefly, MEFs of indicated genotypes were grown in 150mm^2^ culture dishes. Cells were washed with ice-cold PBS, pH 7.4. An equal number of cells were suspended in intracellular buffer (120 mM KCl, 10 mM NaCl, 1 mM KH2PO4, 20 mM HEPES-Tris, pH 7.2) supplemented with SIGMAFAST™ Protease Inhibitor Cocktail (Sigma-Aldrich, Cat#S8830), and permeabilized with digitonin (80 µg/mL) for 5 min at room temperature. The cytochrome c release was induced by adding tBid (20nM) and incubating the cell suspension at 30°C for 30 min. Cell homogenates were spun at 16,500 g, 4°C for 10 min, and the supernatant (cytosolic fraction) was removed. Pellets were lysed 1XRIPA buffer and centrifuged at 16,500 g, 4°C for 10 min to obtain the total cell lysate. Both total cell lysate and cytosolic fractions were immunoblotted using the cytochrome c antibody.

### _m_Ca^2+^ flux analysis

_m_Ca^2+^ flux was analyzed as described earlier (Luongo et al., 2015; Tomar et al., 2016). Briefly, cells were washed in Ca^2+^-free DPBS (Thermo Fisher Scientific, Cat#14190235). An equal number of cells (7×10^6^ cells) were resuspended and permeabilized with 40 µg/ml digitonin in 1.5 ml of intracellular medium (120 mM KCl, 10 mM NaCl, 1 mM KH2PO4, 20 mM HEPES-Tris, pH 7.2), containing 2 µM thapsigargin to block the SERCA pump, and supplemented with 5 mM succinate. Fura-FF (1µM) was loaded to the cell suspension, and fluorescence was monitored in a multiwavelength excitation dual-wavelength emission fluorimeter (Delta RAM, PTI). Extramitochondrial Ca^2+^ is shown as the excitation ratio (340 nm/380 nm) of Fura-FF fluorescence. A Ca^2+^ bolus, and then mitochondrial uncoupler, FCCP (2 µM), were added at the indicated time points (8, 9). All the experiments were performed at 37°C with constant stirring.

### Statistical analysis

Results are presented as mean +/- standard error of the mean. Statistical analysis was performed using GraphPad PRISM 7.05 (Graph Pad Software). All experiments were repeated independently at least three times. Column analyses were performed using an unpaired, 2-tailed t-test (for 2 groups) with Welch’s correction. For grouped analyses, 2-way ANOVA with Tukey post-hoc analysis was performed. *P* values less than 0.05 (95% confidence interval) were considered significant.

### Data availability

All relevant data are available from the authors.

## Acknowledgments

The authors thanks to Trevor Tierney for technical and managerial assistance in the Elrod Laboratory.

## Author Contributions

Conceptualization, D.T. and J.W.E.; Methodology, D.T., M.T., J.F.G., D.W.K., O.S., P.J., A.C.C., and J.W.E.; Investigation, D.T., M.T., O.S., P.J., and J.W.E; Resources, D.T., J.F.G., D.W.K., and J.W.E.; Writing – Original Draft, D.T. and J.W.E; Writing –Review & Editing, D.T., M.T., J.F.G., P.J., and J.W.E.; Supervision, J.W.E.; Funding Acquisition, D.T., and J.W.E.

## Supplementary Figure Legends

**Figure S1:**
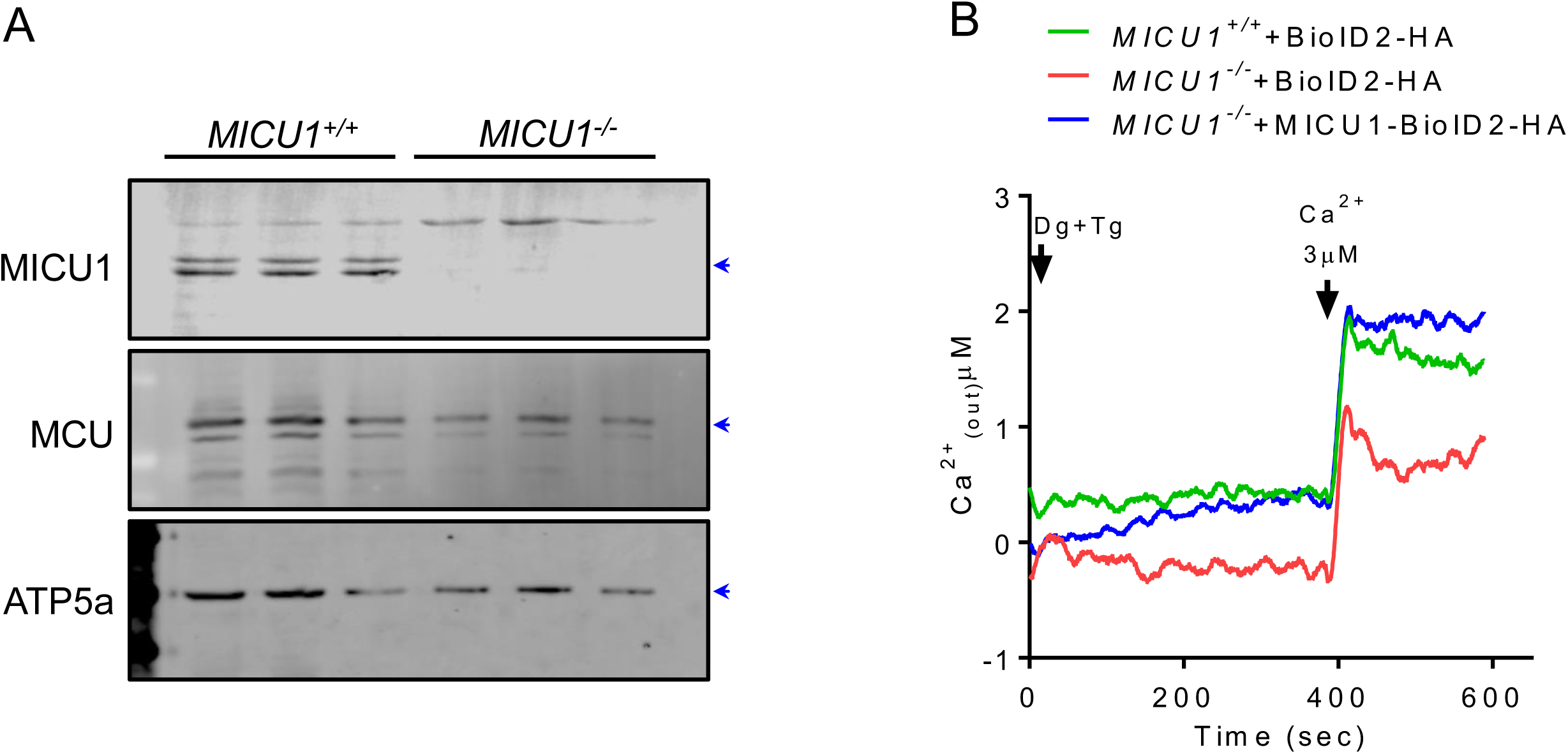
**(A)** Western blot confirming the loss of MICU1 in *Micu1*^-/-^MEFs. **(B)** Representative trace showing _m_Ca^2+^ uptake in WT and *MICU1*^-/-^ HEK293T cells expressing BioID2-HA, or MICU1-BioID2-HA fusion protein.

**Figure S2:**
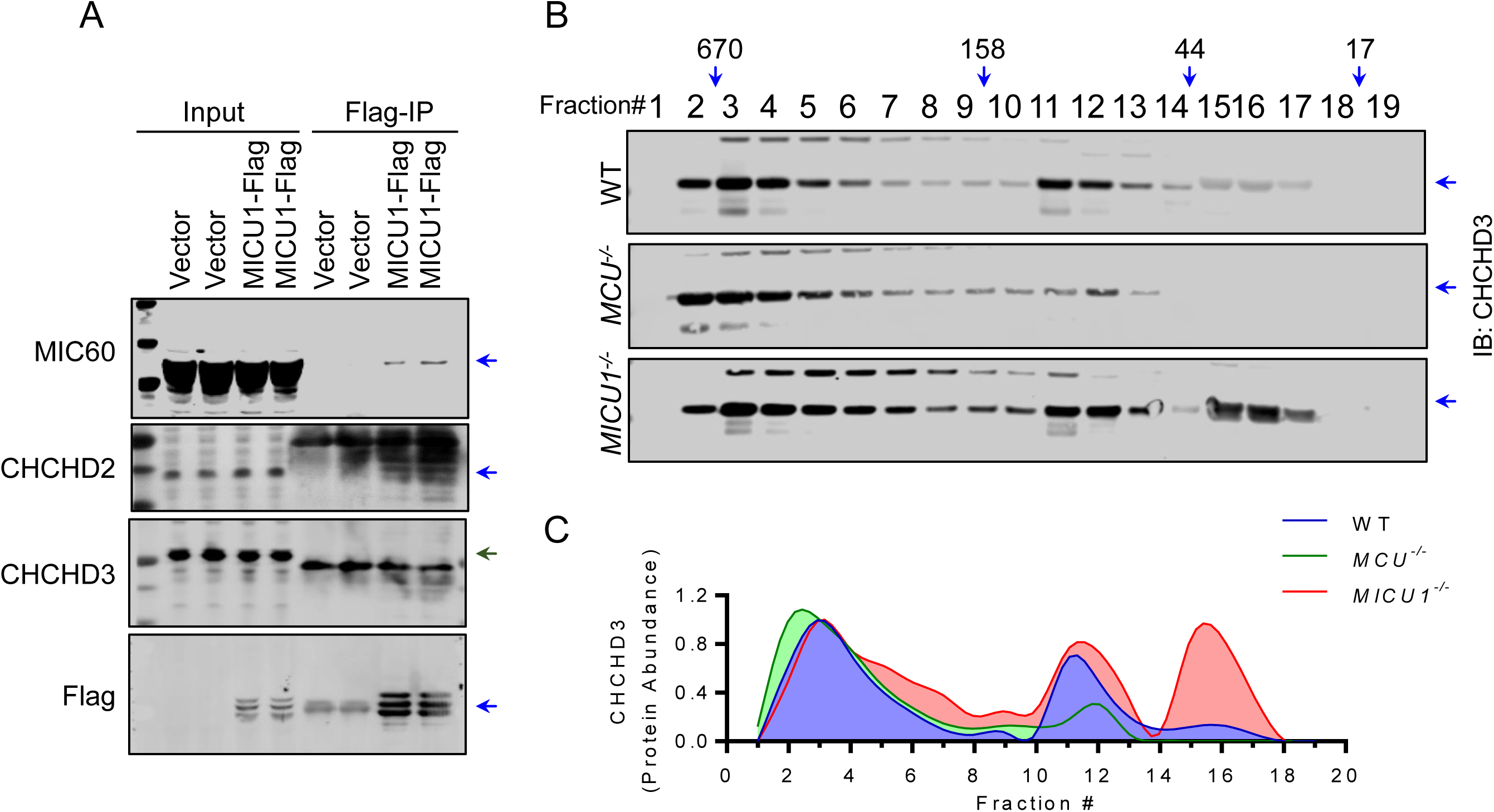
**(A)** MICU1-FLAG was reconstituted in *MICU1*^-/-^ HEK293T cells, and FLAG-IP was performed to detect the interaction with endogenous MICOS components. **(B)** FPLC was performed using cleared cell lysates from WT, *MCU*^-/-^, and *MICU1*^-/-^ HEK293T cells, and immunoblotted for CHCHD3. **(C)** Densitometry was performed to generate chromatographs of the CHCHD3 distribution in different fractions from the cells shown in Figure S2B.

**Figure S3.**
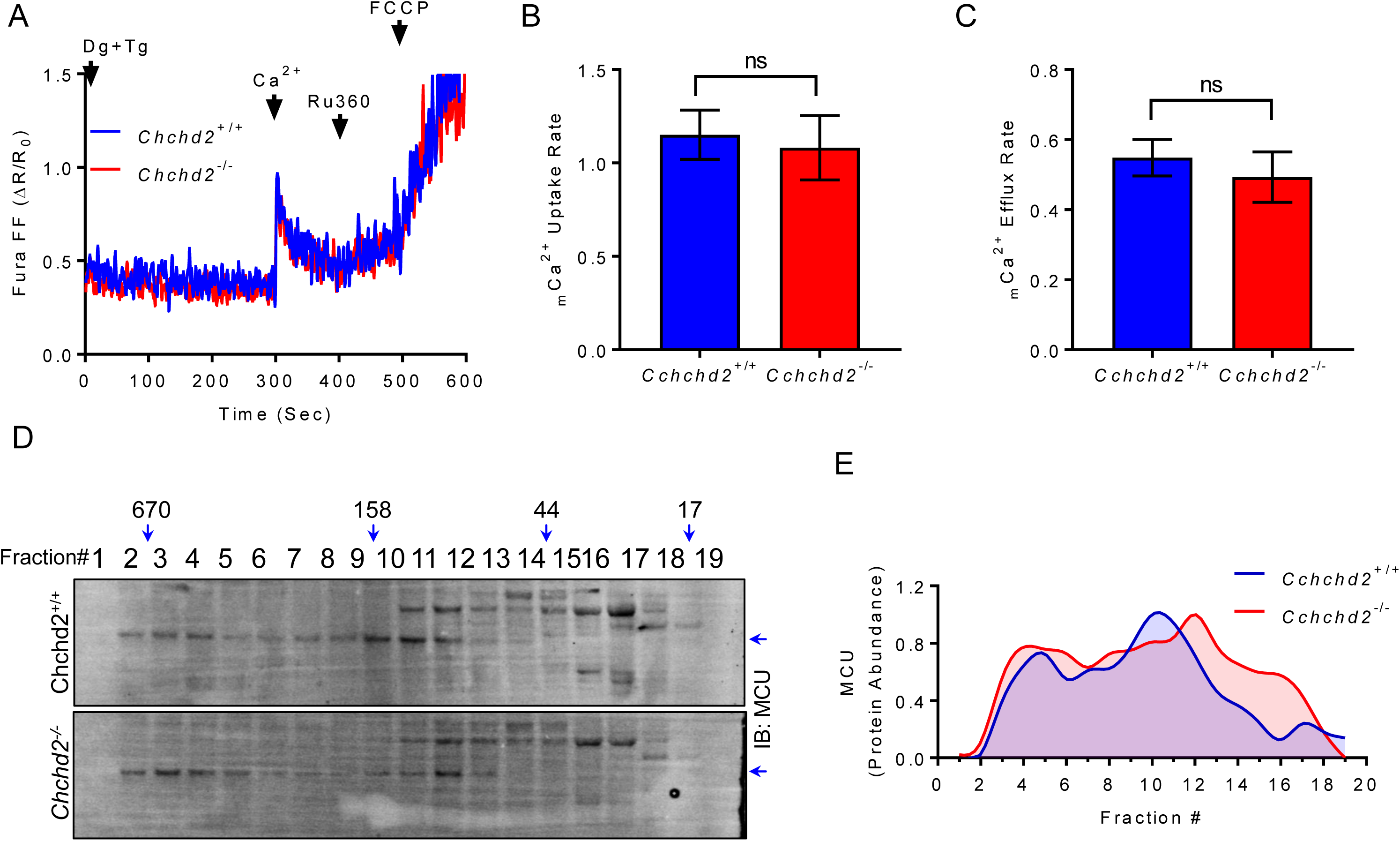
**(A)** Representative trace for _m_Ca^2+^ influx/efflux in *Chchd2*^+/+^ and *Chchd2*^-/-^ MEFs. (**B-C)** _m_Ca^2+^ uptake and efflux rate. **(D-E)** FPLC analysis for native MCU protein complexes distribution in *Chchd2*^+/+^ and *Chchd2*^-/-^ MEFs. The loss of *Chchd2* does not have any effect on the size distribution of the mtCU complex.

**Table S1:** MICU1 proximal MICOS components protein abundance in *MCU*^+/+^ and *MCU*^-/-^ cells

## References

Antony, A.N., Paillard, M., Moffat, C., Juskeviciute, E., Correnti, J., Bolon, B., Rubin, E., Csordas, G., Seifert, E.L., Hoek, J.B., et al. (2016). MICU1 regulation of mitochondrial Ca(2+) uptake dictates survival and tissue regeneration. Nat Commun 7, 10955.

Bagur, R., and Hajnoczky, G. (2017). Intracellular Ca(2+) Sensing: Its Role in Calcium Homeostasis and Signaling. Mol Cell 66, 780–788.

Baughman, J.M., Perocchi, F., Girgis, H.S., Plovanich, M., Belcher-Timme, C.A., Sancak, Y., Bao, X.R., Strittmatter, L., Goldberger, O., Bogorad, R.L., et al. (2011). Integrative genomics identifies MCU as an essential component of the mitochondrial calcium uniporter. Nature 476, 341–345.

Bick, A.G., Wakimoto, H., Kamer, K.J., Sancak, Y., Goldberger, O., Axelsson, A., DeLaughter, D.M., Gorham, J.M., Mootha, V.K., Seidman, J.G., et al. (2017). Cardiovascular homeostasis dependence on MICU2, a regulatory subunit of the mitochondrial calcium uniporter. Proc Natl Acad Sci U S A 114, E9096–E9104.

Carafoli, E. (2002). Calcium signaling: a tale for all seasons. Proc Natl Acad Sci U S A 99, 1115–1122.

Carafoli, E. (2003). The calcium-signalling saga: tap water and protein crystals. Nat Rev Mol Cell Biol 4, 326–332.

Csordas, G., Golenar, T., Seifert, E.L., Kamer, K.J., Sancak, Y., Perocchi, F., Moffat, C., Weaver, D., de la Fuente Perez, S., Bogorad, R., et al. (2013). MICU1 controls both the threshold and cooperative activation of the mitochondrial Ca(2)(+) uniporter. Cell Metab 17, 976–987.

De Stefani, D., Raffaello, A., Teardo, E., Szabo, I., and Rizzuto, R. (2011). A forty-kilodalton protein of the inner membrane is the mitochondrial calcium uniporter. Nature 476, 336–340.

Fransson, A., Ruusala, A., and Aspenstrom, P. (2003). Atypical Rho GTPases have roles in mitochondrial homeostasis and apoptosis. J Biol Chem 278, 6495–6502.

Frederick, R.L., McCaffery, J.M., Cunningham, K.W., Okamoto, K., and Shaw, J.M. (2004). Yeast Miro GTPase, Gem1p, regulates mitochondrial morphology via a novel pathway. J Cell Biol 167, 87–98.

Friedman, J.R., Mourier, A., Yamada, J., McCaffery, J.M., and Nunnari, J. (2015). MICOS coordinates with respiratory complexes and lipids to establish mitochondrial inner membrane architecture. Elife 4.

Gottschalk, B., Klec, C., Waldeck-Weiermair, M., Malli, R., and Graier, W.F. (2018). Intracellular Ca(2+) release decelerates mitochondrial cristae dynamics within the junctions to the endoplasmic reticulum. Pflugers Arch 470, 1193–1203.

Greenawalt, J.W., Rossi, C.S., and Lehninger, A.L. (1964). Effect of Active Accumulation of Calcium and Phosphate Ions on the Structure of Rat Liver Mitochondria. J Cell Biol 23, 21–38.

Harner, M., Korner, C., Walther, D., Mokranjac, D., Kaesmacher, J., Welsch, U., Griffith, J., Mann, M., Reggiori, F., and Neupert, W. (2011). The mitochondrial contact site complex, a determinant of mitochondrial architecture. EMBO J 30, 4356–4370.

Hoffman, N.E., Chandramoorthy, H.C., Shamugapriya, S., Zhang, X., Rajan, S., Mallilankaraman, K., Gandhirajan, R.K., Vagnozzi, R.J., Ferrer, L.M., Sreekrishnanilayam, K., et al. (2013). MICU1 motifs define mitochondrial calcium uniporter binding and activity. Cell Rep 5, 1576–1588.

Kim, D.I., Jensen, S.C., Noble, K.A., Kc, B., Roux, K.H., Motamedchaboki, K., and Roux, K.J. (2016). An improved smaller biotin ligase for BioID proximity labeling. Mol Biol Cell 27, 1188–1196.

Kirichok, Y., Krapivinsky, G., and Clapham, D.E. (2004). The mitochondrial calcium uniporter is a highly selective ion channel. Nature 427, 360–364.

Lewis-Smith, D., Kamer, K.J., Griffin, H., Childs, A.M., Pysden, K., Titov, D., Duff, J., Pyle, A., Taylor, R.W., Yu-Wai-Man, P., et al. (2016). Homozygous deletion in MICU1 presenting with fatigue and lethargy in childhood. Neurol Genet 2, e59.

Liu, J.C., Liu, J., Holmstrom, K.M., Menazza, S., Parks, R.J., Fergusson, M.M., Yu, Z.X., Springer, D.A., Halsey, C., Liu, C., et al. (2016). MICU1 Serves as a Molecular Gatekeeper to Prevent In Vivo Mitochondrial Calcium Overload. Cell Rep 16, 1561–1573.

Logan, C.V., Szabadkai, G., Sharpe, J.A., Parry, D.A., Torelli, S., Childs, A.M., Kriek, M., Phadke, R., Johnson, C.A., Roberts, N.Y., et al. (2014). Loss-of-function mutations in MICU1 cause a brain and muscle disorder linked to primary alterations in mitochondrial calcium signaling. Nat Genet 46, 188–193.

Luongo, T.S., Lambert, J.P., Yuan, A., Zhang, X., Gross, P., Song, J., Shanmughapriya, S., Gao, E., Jain, M., Houser, S.R., et al. (2015). The Mitochondrial Calcium Uniporter Matches Energetic Supply with Cardiac Workload during Stress and Modulates Permeability Transition. Cell Rep 12, 23–34.

Mallilankaraman, K., Cardenas, C., Doonan, P.J., Chandramoorthy, H.C., Irrinki, K.M., Golenar, T., Csordas, G., Madireddi, P., Yang, J., Muller, M., et al. (2012a). MCUR1 is an essential component of mitochondrial Ca2+ uptake that regulates cellular metabolism. Nat Cell Biol 14, 1336–1343.

Mallilankaraman, K., Doonan, P., Cardenas, C., Chandramoorthy, H.C., Muller, M., Miller, R., Hoffman, N.E., Gandhirajan, R.K., Molgo, J., Birnbaum, M.J., et al. (2012b). MICU1 is an essential gatekeeper for MCU-mediated mitochondrial Ca(2+) uptake that regulates cell survival. Cell 151, 630–644.

Meng, H., Yamashita, C., Shiba-Fukushima, K., Inoshita, T., Funayama, M., Sato, S., Hatta, T., Natsume, T., Umitsu, M., Takagi, J., et al. (2017). Loss of Parkinson’s disease-associated protein CHCHD2 affects mitochondrial crista structure and destabilizes cytochrome c. Nat Commun 8, 15500.

Nemani, N., Carvalho, E., Tomar, D., Dong, Z., Ketschek, A., Breves, S.L., Jana, F., Worth, A.M., Heffler, J., Palaniappan, P., et al. (2018). MIRO-1 Determines Mitochondrial Shape Transition upon GPCR Activation and Ca(2+) Stress. Cell Rep 23, 1005–1019.

Paillard, M., Csordas, G., Huang, K.T., Varnai, P., Joseph, S.K., and Hajnoczky, G. (2018). MICU1 Interacts with the D-Ring of the MCU Pore to Control Its Ca(2+) Flux and Sensitivity to Ru360. Mol Cell 72, 778–785 e773.

Patron, M., Checchetto, V., Raffaello, A., Teardo, E., Vecellio Reane, D., Mantoan, M., Granatiero, V., Szabo, I., De Stefani, D., and Rizzuto, R. (2014). MICU1 and MICU2 finely tune the mitochondrial Ca2+ uniporter by exerting opposite effects on MCU activity. Mol Cell 53, 726–737.

Perocchi, F., Gohil, V.M., Girgis, H.S., Bao, X.R., McCombs, J.E., Palmer, A.E., and Mootha, V.K. (2010). MICU1 encodes a mitochondrial EF hand protein required for Ca(2+) uptake. Nature 467, 291–296.

Phillips, C.B., Tsai, C.W., and Tsai, M.F. (2019). The conserved aspartate ring of MCU mediates MICU1 binding and regulation in the mitochondrial calcium uniporter complex. Elife 8.

Plovanich, M., Bogorad, R.L., Sancak, Y., Kamer, K.J., Strittmatter, L., Li, A.A., Girgis, H.S., Kuchimanchi, S., De Groot, J., Speciner, L., et al. (2013). MICU2, a paralog of MICU1, resides within the mitochondrial uniporter complex to regulate calcium handling. PLoS One 8, e55785.

Raffaello, A., De Stefani, D., Sabbadin, D., Teardo, E., Merli, G., Picard, A., Checchetto, V., Moro, S., Szabo, I., and Rizzuto, R. (2013). The mitochondrial calcium uniporter is a multimer that can include a dominant-negative pore-forming subunit. EMBO J 32, 2362–2376.

Rizzuto, R., De Stefani, D., Raffaello, A., and Mammucari, C. (2012). Mitochondria as sensors and regulators of calcium signalling. Nat Rev Mol Cell Biol 13, 566–578.

Rizzuto, R., and Pozzan, T. (2006). Microdomains of intracellular Ca2+: molecular determinants and functional consequences. Physiol Rev 86, 369–408.

Sancak, Y., Markhard, A.L., Kitami, T., Kovacs-Bogdan, E., Kamer, K.J., Udeshi, N.D., Carr, S.A., Chaudhuri, D., Clapham, D.E., Li, A.A., et al. (2013). EMRE is an essential component of the mitochondrial calcium uniporter complex. Science 342, 1379–1382.

Saotome, M., Safiulina, D., Szabadkai, G., Das, S., Fransson, A., Aspenstrom, P., Rizzuto, R., and Hajnoczky, G. (2008). Bidirectional Ca2+-dependent control of mitochondrial dynamics by the Miro GTPase. Proc Natl Acad Sci U S A 105, 20728–20733.

Scorrano, L., Ashiya, M., Buttle, K., Weiler, S., Oakes, S.A., Mannella, C.A., and Korsmeyer, S.J. (2002). A distinct pathway remodels mitochondrial cristae and mobilizes cytochrome c during apoptosis. Dev Cell 2, 55–67.

Tadross, M.R., Dick, I.E., and Yue, D.T. (2008). Mechanism of local and global Ca2+ sensing by calmodulin in complex with a Ca2+ channel. Cell 133, 1228–1240.

Tarasenko, D., Barbot, M., Jans, D.C., Kroppen, B., Sadowski, B., Heim, G., Mobius, W., Jakobs, S., and Meinecke, M. (2017). The MICOS component Mic60 displays a conserved membrane-bending activity that is necessary for normal cristae morphology. J Cell Biol 216, 889–899.

Tomar, D., Dong, Z., Shanmughapriya, S., Koch, D.A., Thomas, T., Hoffman, N.E., Timbalia, S.A., Goldman, S.J., Breves, S.L., Corbally, D.P., et al. (2016). MCUR1 Is a Scaffold Factor for the MCU Complex Function and Promotes Mitochondrial Bioenergetics. Cell Rep 15, 1673–1685.

Tomar, D., Prajapati, P., Lavie, J., Singh, K., Lakshmi, S., Bhatelia, K., Roy, M., Singh, R., Benard, G., and Singh, R. (2015). TRIM4; a novel mitochondrial interacting RING E3 ligase, sensitizes the cells to hydrogen peroxide (H2O2) induced cell death. Free Radic Biol Med 89, 1036–1048.

Tufi, R., Gleeson, T.P., von Stockum, S., Hewitt, V.L., Lee, J.J., Terriente-Felix, A., Sanchez-Martinez, A., Ziviani, E., and Whitworth, A.J. (2019). Comprehensive Genetic Characterization of Mitochondrial Ca(2+) Uniporter Components Reveals Their Different Physiological Requirements In Vivo. Cell Rep 27, 1541–1550 e1545.

van der Laan, M., Horvath, S.E., and Pfanner, N. (2016). Mitochondrial contact site and cristae organizing system. Curr Opin Cell Biol 41, 33–42.

Xing, Y., Wang, M., Wang, J., Nie, Z., Wu, G., Yang, X., and Shen, Y. (2019). Dimerization of MICU Proteins Controls Ca(2+) Influx through the Mitochondrial Ca(2+) Uniporter. Cell Rep 26, 1203–1212 e1204.

